# Curating models from BioModels: Developing a workflow for creating OMEX files

**DOI:** 10.1101/2024.03.15.585236

**Authors:** Jin Xu, Lucian Smith

**Affiliations:** Department of Bioengineering, University of Washington, Seattle, WA, USA

## Abstract

The reproducibility of computational biology models can be greatly facilitated by widely adopted standards and public repositories. We examined 50 models from the BioModels Database and attempted to validate the original curation and correct some of them if necessary. For each model, we reproduced these published results using Tellurium. Once reproduced we manually created a new set of files, with the model information stored by the Systems Biology Markup Language (SBML), and simulation instructions stored by the Simulation Experiment Description Markup Language (SED-ML), and everything included in an Open Modeling EXchange (OMEX) file, which could be used with a variety of simulators to reproduce the same results. On the one hand, the reproducibility procedure of 50 models developed a manual workflow that we would use to build an automatic platform to help users more easily curate and verify models in the future. On the other hand, these exercises allowed us to find the limitations and possible enhancement of the current curation and tooling to verify and curate models.

## Introduction

Because discoveries are almost always dependent on previous results, methodologies, and theories, reproducibility has become a fundamental part of the scientific process [1]. Reproducibility of methods requires one to be able to exactly reproduce the results using the same methods on the same data, while reproducibility of results requires one to obtain similar results in an independent study applying similar procedures [2].

Therefore, the deposition of models in public repositories using standard formats like the Systems Biology Markup Language (SBML) [3] or CellML [4] has been an important resource in computational systems biology. The repositories allow scientists to easily find and access models, use them to run simulations and derive new models and simulations using compatible software applications. During the last couple of decades, many classic models have been added to model repositories. Public standards and repositories can facilitate the reuse and regeneration of computational biology models that will outlive the original used specific software [5].

The BioModels Database (https://www.ebi.ac.uk/biomodels/) [6, 7] is one of the largest public open-source databases for quantitative biological models, where the models are manually curated and enriched. The curation includes but is not limited to the validity of the model file and whether the model provides results corresponding to the reference scientific article [6]. However, there are some limitations of the current curation efforts for the BioModels Database. For instance, some curated plots are not the same as those found in the published papers, and the description of how the plots were created is limited to a text listing. Storing this information using the Simulation Experiment Description Markup Language (SED-ML) [8] has the potential to encode these experiments and to be extended to cover more results from the paper. SED-ML files are present in about a third of the BioModels Database (373 of 1058 entries), but have not been validated nor verified against the curated plots.

To validate and correct the curated models, we examined 50 models from the BioModels Database, and successfully reproduced published results using Tellurium [9, 10]. Once reproduced we updated the existing SED-ML file or created a new SED-ML file which repeated this experiment, and stored this with the original SBML model in an Open Modeling EXchange (OMEX) [11] file. OMEX is the basis of the Computational Modeling in Biological Network (COMBINE) Archive, with a single file supporting the exchange of all the information necessary for a modeling and simulation experiment in biology [12]. The input to each tool of BioSimulations (https://biosimulations.org) is a COMBINE archive which contains SED-ML files that describe simulations of models in formats such as SBML [13]. The tools from BioSimulations include but are not limited to Tellurium, COPASI [14], and VCell [15], etc.

The successful reproduction of the 50 models suggested a certain manual workflow to generate OMEX files. During the reproducing process, some curated results were corrected and extended. However, only 19 among the 50 models of our curation covered all the model-related figures in the corresponding papers, which means there were still many results that were not covered. Therefore, our work also identified some issues in reproducing models from the perspective of tooling and papers to achieve reproducibility. We found some limitations in the tooling and papers to achieve reproducibility and suggested some possible enhancements for curation and tooling in the future.

## Materials and methods

There are over a thousand models available in the BioModels Database. To demonstrate how to validate, correct, and extend the current curation of BioModels entries, we examined a selection of models to develop a manual workflow to generate OMEX files. Here, we present a systematic analysis of model reproducibility by attempting to independently reproduce published modeling results. In total, we investigated 50 models selected from the BioModels Database. Initially, we selected the 50 models because they seemed easy to reproduce, get fixed, or extended. However, we found the 50 models could represent some current issues regarding reproducibility and curation in the BioModels Database and tooling sets. First of all, not all the models include SED-ML files, so it is difficult to reproduce the curation. Secondly, some of the SED-ML files do not produce plots that match the corresponding figures in the paper. Thirdly, there are usually multiple figures in the paper, however, the current curation only provides some reproduced figures instead of all of them. In this work, we manually validated or corrected and extended the SED-ML for all 50 entries. Following this procedure, we also found the limitation to reproduce or extend the current curation due to tooling and the paper information.

In BioModels Database, the website of each model provides several sections including “Overview”, “Files”, and “Curation”. We made use of the “Format Related Publication” in the “Overview” section to download the corresponding article to access detailed information from the paper to reproduce its results. In the “Files” section, there is the SBML file providing the model information and there are sometimes SED-ML files providing simulation information that we used. In the “Curation” section, there are plots curated by BioModels with which we compared our own generated plots.

We reproduced the published results using Tellurium version 2.2.5.2 that imports RoadRunner version 2.3.1 [16, 17]. Once reproduced, we manually created a standard OMEX file using SBML and SED-ML following the manual workflow steps below.

### Read and modify the SBML file

To reproduce figures other than the one initially curated, values in the model itself can often be changed. This can be accomplished manually using tools such as Antimony [18] and libSBML [19]. We can also modify SBML files manually and directly.

For example, the curation for BIOMD0000000720 [20] (https://www.ebi.ac.uk/biomodels/BIOMD0000000720#Curation) only provides the reproduced figure Fig 7a. We were able to extend this to reproduce Figures 6, 7, and 9 by adjusting the appropriate parameters and initial values, as listed in Table 1. In Fig 1, we have shown the successfully reproduced Fig 7a and extended Fig 7b and Fig 9 as representations. The parameter values were available in the SBML files. In addition, we referred to the papers for parameters to validate, correct, and extend the curations. See S1 Table for details about finding parameters in papers.

**Table 1.** The 50 reproduced models from BioModels Database.

| BioModel ID | Paper | Figure | Improvement | BioModel ID | Paper | Figure | Improvement |
| --- | --- | --- | --- | --- | --- | --- | --- |
| BIOMD0000000003 | [21] | Fig 3 | validated | BIOMD0000000850 | [22] | Fig 3-1, 3-2 | extended |
| BIOMD0000000005 | [23] | Fig 3a | validated | BIOMD0000000877 | [24] | Fig 1, 2 | extended |
| BIOMD0000000010 | [25] | Fig 2A | corrected | BIOMD0000000894 | [26] | Fig 3a | validated |
| BIOMD0000000079 | [27] | Fig 3, 4, 6 | extended | BIOMD0000000909 | [28] | Fig 10 | extended |
| BIOMD0000000548 | [29] | Fig 3 | validated | BIOMD0000000911 | [30] | Fig 1 | validated |
| BIOMD0000000552 | [31] | Fig 1 | validated | BIOMD0000000916 | [32] | Fig 2, 3 | corrected |
| BIOMD0000000555 | [33] | Fig 1 | validated | BIOMD0000000930 | [34] | Fig 1 | extended |
| BIOMD0000000618 | [35] | Fig 4 | validated | BIOMD0000000932 | [36] | Fig 4 | validated |
| BIOMD0000000642 | [37] | Fig 1 | validated | BIOMD0000000933 | [38] | Fig 3 | validated |
| BIOMD0000000667 | [39] | Fig 2 | validated | BIOMD0000000939 | [40] | Fig 2-7 | extended |
| BIOMD0000000671 | [41] | Fig 3 | validated | BIOMD0000000947 | [42] | Fig 6 | validated |
| BIOMD0000000704 | [43] | Fig 8, 9 | extended | BIOMD0000000948 | [44] | Fig 3 | validated |
| BIOMD0000000712 | [45] | Fig 2 | extended | BIOMD0000000949 | [46] | Fig 2 | validated |
| BIOMD0000000720 | [20] | Fig 6, 7, 9 | extended | BIOMD0000000953 | [47] | Fig 7B | extended |
| BIOMD0000000745 | [48] | Fig 6 up | extended | BIOMD0000000964 | [49] | Fig 2 | corrected |
| BIOMD0000000757 | [50] | Fig 1 | validated | BIOMD0000000967 | [51] | Fig 2 | validated |
| BIOMD0000000780 | [52] | Fig 6 | validated | BIOMD0000000970 | [53] | Fig 2 | corrected |
| BIOMD0000000781 | [52] | Fig 4 | validated | BIOMD0000000984 | [54] | Fig 3 | validated |
| BIOMD0000000782 | [52] | Fig 1-3 | validated | BIOMD0000000986 | [55] | Fig 2 | validated |
| BIOMD0000000785 | [56] | Fig 3b | validated | BIOMD0000001004 | [57] | Fig 3 | validated |
| BIOMD0000000793 | [58] | Fig 2, 3 | validated | BIOMD0000001006 | [59] | Fig 2 | corrected |
| BIOMD0000000795 | [58] | Fig 4 | validated | BIOMD0000001023 | [60] | Fig 5 | validated |
| BIOMD0000000799 | [61] | Fig 8-10 | extended | BIOMD0000001026 | [62] | Fig 7 | extended |
| BIOMD0000000815 | [63] | Fig 5-7 | extended | BIOMD0000001037 | [64] | Fig 10 | validated |
| BIOMD0000000839 | [65] | Fig 2, 3 | corrected | BIOMD0000001038 | [64] | Fig 11 | validated |

The BioModel ID, Paper, Figure, and Improvement columns are for the model IDs in the BioModels Database, the published journal articles where the models were originally from, the reproduced figure indices in the paper correspondingly, and our contribution to improve the original curation. “Validated” means that the original curated figure is correct, and we successfully reproduced the curated result and did not reproduce additional results from the original article. “Corrected” means that the original curated figure is incorrect, but we successfully corrected the results to be the same as in the original article. “Extended” means that the original curated figure is correct, but we reproduced more results beyond the original curation.

**Fig 1.**
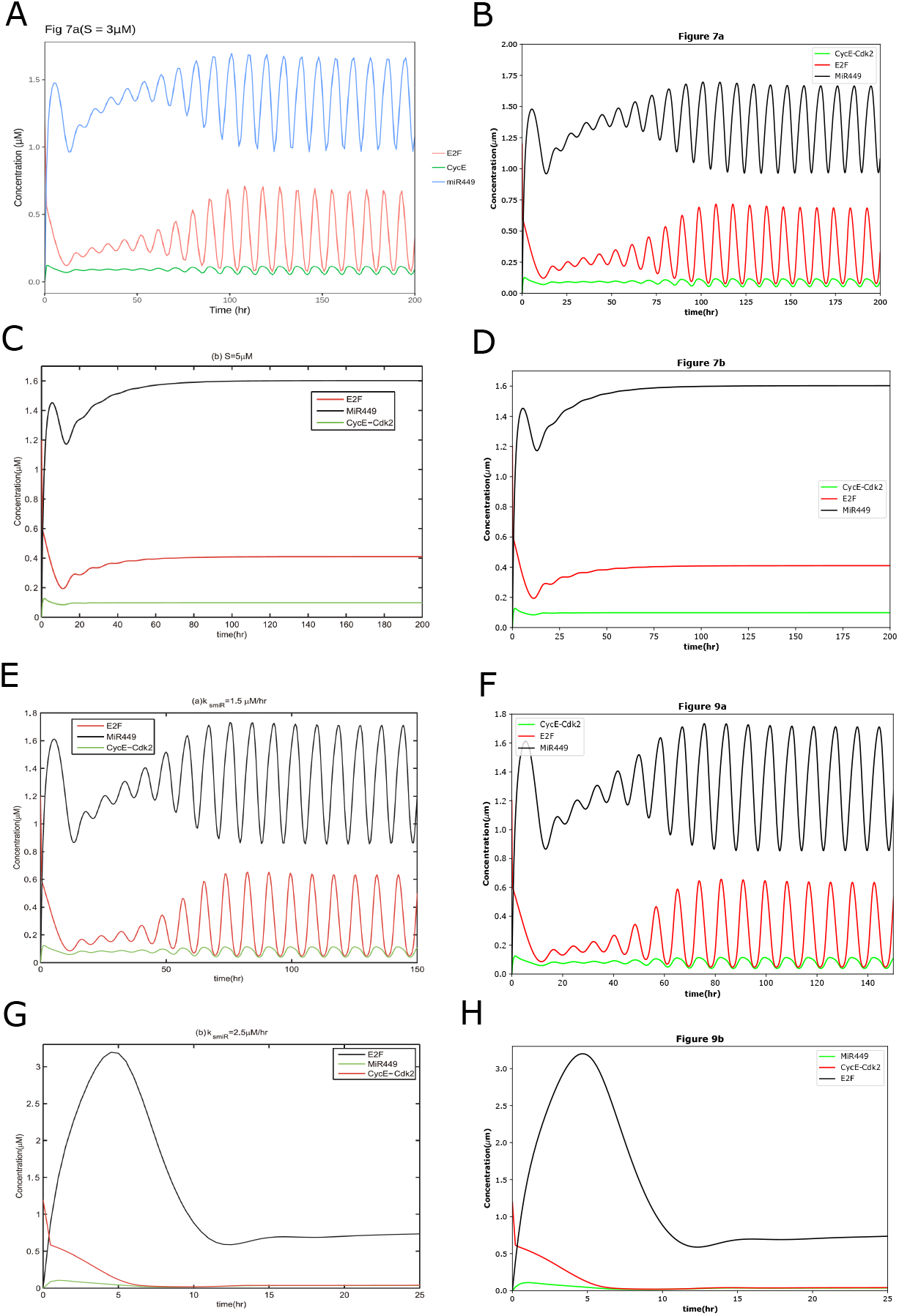
The reproduced results based on BIOMD0000000720. BIOMD0000000720 describes the dynamical behaviors of Rb-E2F pathway including negative feedback loops involving miR449. (A) is the original curation from the BioModels Database which is comparable with Fig 7a in the original paper [20] except line styles. (C), (E), and (G) are the original results published in the paper as Fig 7b and Fig 9. The figures illustrate the time courses of [E2F], [CycE-Cdk2] and [MiR449] with different values of *k*_*smiR*_ and *S*. (B), (D), (F), and (H) indicated the comparable results reproduced by Tellurium.

### Read and modify the SED-ML file

SED-ML is a representation format based on XML for the encoding and exchange of simulation descriptions on computational models of biological systems. It stores all the simulation information of a certain biology model. We first read the simulation information of the model from the SED-ML file. The model simulation information includes but is not limited to the model to simulate time courses and output formats. Some curation might not be correct, then we needed to correct the curation by adjusting the SED-ML files. We used phraSED-ML [66] and libSEDML [67] to achieve the modification.

For example, the curation of Figure 3 for BIOMD0000000916 (https://www.ebi.ac.uk/biomodels/BIOMD0000000916#Curation) is incorrect, with saturation around 0.8 instead of 1. We noticed that the original curation only considered the contribution from *X*_2_ to make the saturation value smaller than in the paper. Therefore, we adjusted the contribution from both *X*_2_ and *X*_3_, and successfully corrected the original curation of Fig 3. In Fig 2, we have shown the corrected figure (Fig 2B) compared with the figure in the paper (Fig 2A) and the original curation (Fig 2C). It is also listed in Table 1.

**Fig 2.**
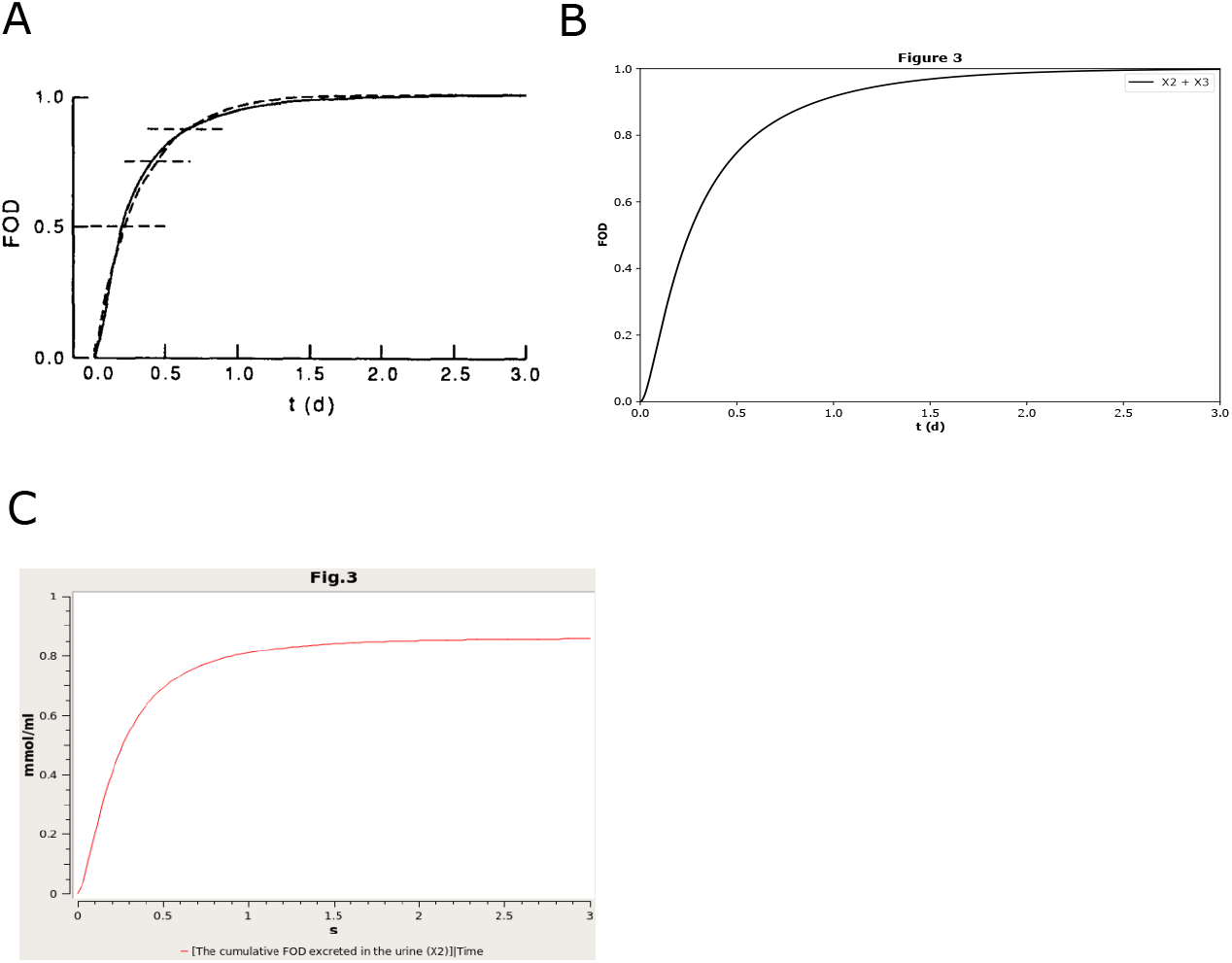
The reproduced results based on BIOMD00000000916. BIOMD0000000916 describes a hypothetic model about the kinetics of control metabolism and excretion. (A) is the original results published in the paper [32] as Fig 3. The figures illustrate the total excretion of the [^3^H]F metabolites from the body as time goes. (B) indicated the comparable results reproduced by Tellurium. (C) is the original curation from the BioModels Database which is not the same as in the corresponding paper.

**Fig 3.**
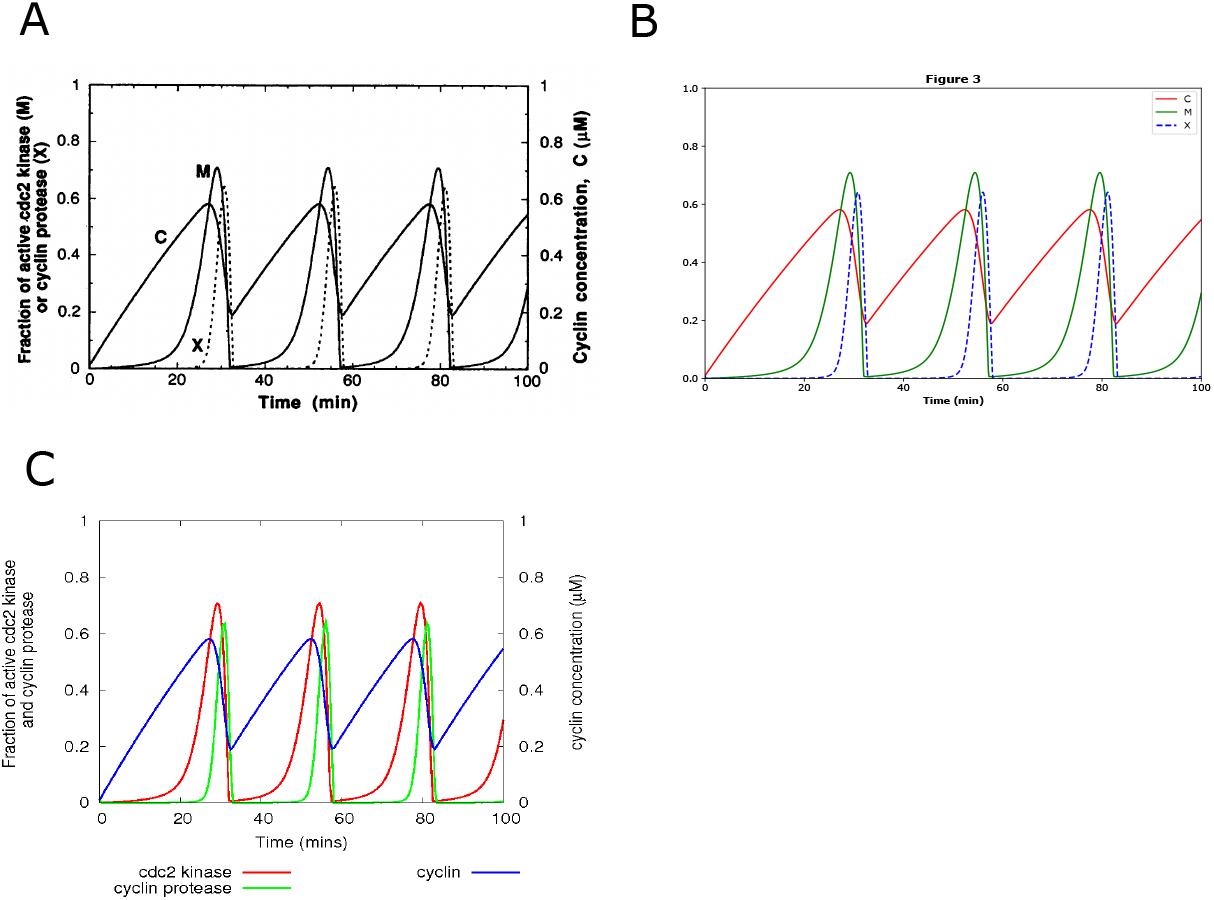
The reproduced results based on BIOMD0000000003. BIOMD0000000003 describes a minimal cascade model for the mitotic oscillator involving cyclin and cdc2 kinase. (A) is the original result published in the paper [21] as Fig 3. The figure shows how the fraction of active cdc2 kinase (M), cyclin protease (X), and cyclin concentration (C) go with time in minutes. (B) indicated the comparable result reproduced by Tellurium. (C) is the original curation from the BioModels Database with comparable results except line styles.

Here, our workflow just assumed that there were SED-ML files that existed and might need modifications. However, some models in BioModels do not provide SED-ML files but only provide SBML model files. Of the 50 models we selected for curation, nine had no existing SED-ML. Therefore, we used some SED-ML files referring to https://github.com/sys-bio/temp-biomodels/tree/main/final.

### Generate figures by libSEDML and create OMEX files

The final steps were to generate comparable plots by Tellurium as the plots shown in the original articles. To make the visualizations vivid, article authors usually use different colors and line styles to represent their results. libSEDML can modify and store the styles of the plots in the SED-ML files.

For example, Fig 3 in the curation for BIOMD0000000003 (https://www.ebi.ac.uk/biomodels/BIOMD0000000003#Curation) had different line styles from the original published article. Then, we adjusted the line style of cyclin protease (X) to dashed lines from solid line style and added some colors to distinguish cdc2 kinase (M) and cyclin concentration (C). In Fig 3, we have shown our reproduced figure with dashed lines (Fig 3B) compared with the figure in the paper (Fig 3A) and the original curation (Fig 3C) without dashed lines. It is also listed in Table 1.

Once we had all the information of the model and the simulation information with its output styles stored in SBML and SED-ML files, we manually created the OMEX files in the end.

The three steps mentioned above allowed us to achieve a manual workflow to generate OMEX files programming by Tellurium in Python. There are two sample scripts about the generation of OMEX files available on GitHub (https://github.com/sys-bio/Developing-a-workflow-for-creating-OMEX-files) under the folder of script examples. The generation process of BIOMD0000000010 was based on the phraSED-ML string, while the generation process of BIOMD0000000003 was based on the SED-ML file. For novice users, we recommend the Windows Installer to install Tellurium with the Spyder Integrated Development Environment (IDE), which is made up of some core building blocks including an “Editor”, an “IPython Console”, “Plots” etc. Users would only need to open the provided file create omex.py in the “Editor” and run it, then could see the generated plots in the “Plots”, and obtain the generated plots, SED-ML, and OMEX files within the same folder of the Python script. The Python scripts, standard files, and generated plots for each model were provided on GitHub under the folder omex of each BioModel. Under the folder of each BioModel, there was the folder paper to provide the original manuscript with parameter information highlighted, the folder original curation to provide the original BioModels curation to compare with, and the folder old SEDML with SED-ML files, if any, before our modification.

## Results

### Successful reproduction with the workflow

We successfully validated, corrected, and extended 50 models from the BioModels Database following the workflow stated in the section Materials and Methods. Table 1 indicates all the reproduced models and their corresponding papers and figures. Among all the 50 models, we validated 30 models, corrected six models, and extended 14 models. “Validated” means that the original curated figure is correct, and we successfully reproduced the curated result and did not reproduce additional results from the original article. “Corrected” means that the original curated figure is incorrect, but we successfully corrected the results to be the same as in the original article. “Extended” means that the original curated figure is correct, but we reproduced more results beyond the original curation. The “Corrected” also included the cases with both correction and extension. We also adjusted the line colors and styles according to the original papers, which were not indicated in Table 1. Among the 50 reproduced models, we adjusted eight models with their line colors and styles to be comparable with the original articles. Here we selected some interesting models as representations to illustrate the current curated BioModels status corresponding to their original published articles.

The first example is a model of “Validated”. There is usually one plot in one paper corresponding to a certain BioModel curation. For example, BIOMD0000001023 (https://www.ebi.ac.uk/biomodels/BIOMD0000001023#Curation) curates the Fig 5 in the corresponding paper [60], which is correct. Here we validated the curation in Tellurium as shown in Fig 4, as an example to illustrate that the manual workflow worked.

**Fig 4.**
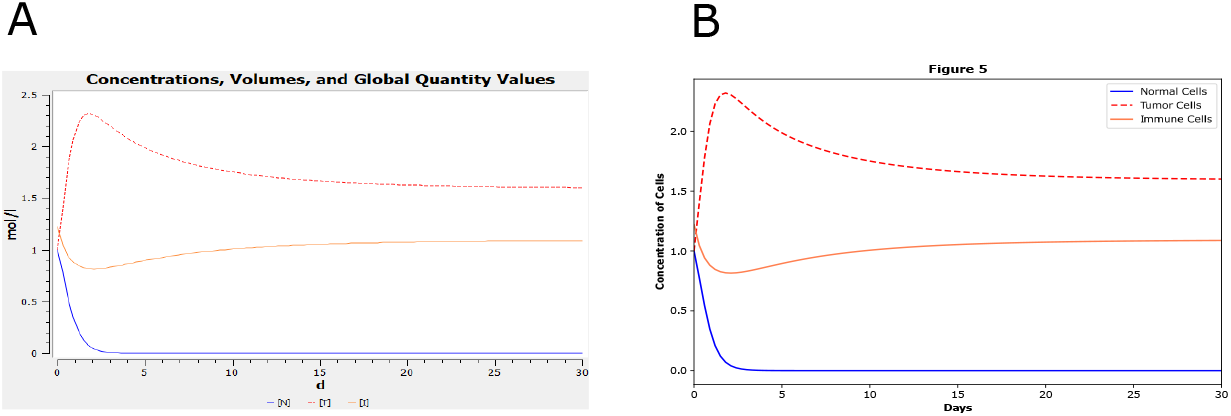
The reproduced results based on BIOMD0000001023. BIOMD0000001023 describes a new ODE-based model for tumor cells and immune system competition. (A) is the original curation from the BioModels Database which is comparable with Fig 5 in the corresponding paper [60]. The figure shows how the concentration of normal, tumor, and immune cells go with time in days. (B) indicated the comparable result validated by Tellurium.

**Fig 5.**
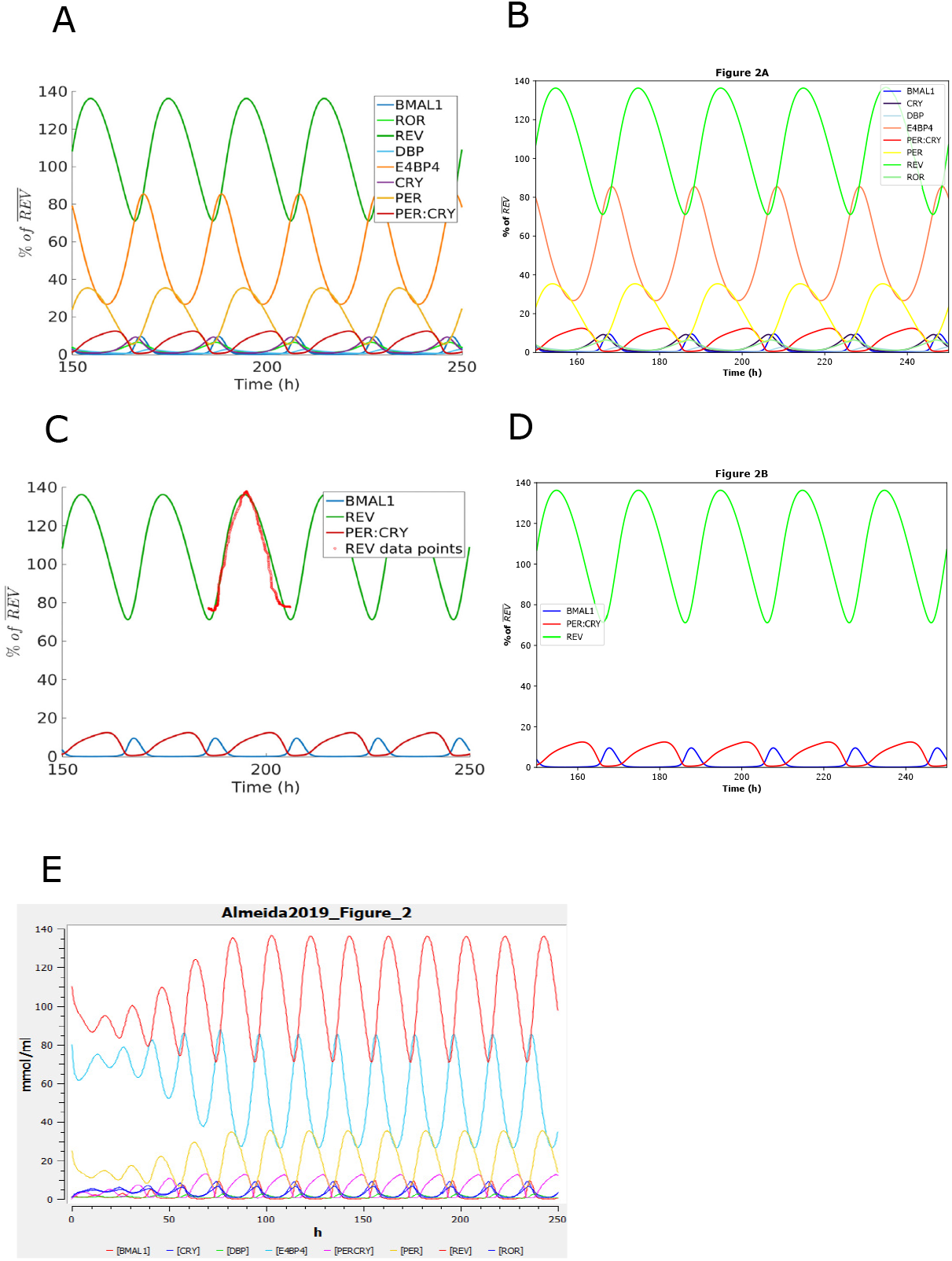
The reproduced results based on BIOMD0000000839. BIOMD0000000839 describes the transcription-based circadian mechanism that controls the duration of molecular clock states in response to signaling inputs. (A) and (C) are the original results published in the paper [65] as Fig 2. The figures illustrate the mammalian circadian clock described by a model focused on transcriptional regulation. (B) and (D) indicate the comparable results reproduced by Tellurium. (E) is the original curation which is not exactly the same as the original paper.

The second example is a model of “Corrected”. Some entries contained extra information in their plots than were present in the published figures, making visual comparison difficult. For example, the entry BIOMD0000000839 (https://www.ebi.ac.uk/biomodels/BIOMD0000000839#Curation) contains the entire simulation from time zero to time 250, while the paper only displays the plot between time points 150 to 250. There is also a time shift in the curated plot compared with the original article. We corrected these plots as shown in Fig 5.

The third example is a model of “Extended”. The model in a BioModels entry can correspond to multiple reproducible plots. In many cases, some plots were reproduced during curation, such as BIOMD0000000939 (https://www.ebi.ac.uk/biomodels/BIOMD0000000939#Curation). In this BioModels entry, the plots from Figures 2, 3, 4, and 6 were reproduced. However, we were able to additionally reproduce more plots from Figures 5 and 7 by adjusting the parameters. Fig 6 provided the validated curation of Fig 4 and Fig 6 in the paper [40] and represented the extra results of Fig 5 and Fig 7 as an extension of the curation.

**Fig 6.**
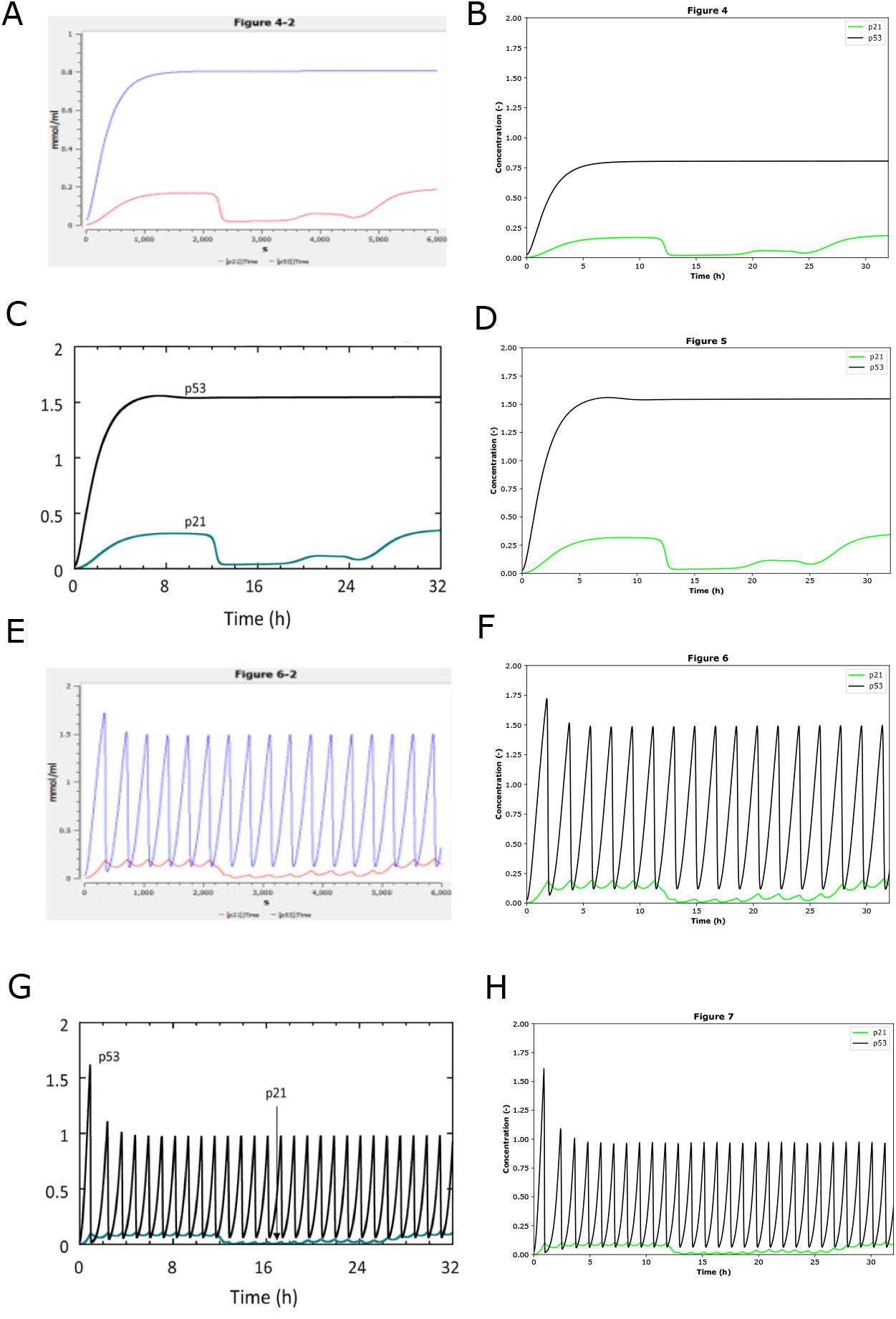
The reproduced results based on BIOMD0000000939. BIOMD0000000939 describes the mathematical modeling of cell cycle regulation in response to DNA damage. (A) and (E) are the original curation which is comparable with Figs 4 and 6 in the paper [40] except the line styles and axis scales. (C) and (G) are the original results published in Figs 5 and 7 in the paper. Figs (A), (C), (E), and (G) illustrate how the concentrations of p53 and p21 go with time in hours with the DNA damage signal (DDS) as 0.002, 0.004, 0.008, 0.016. (B), (D), (F), and (H) indicated the comparable results reproduced by Tellurium.

In some cases, there are multiple BioModels corresponding to one single paper. Therefore, a possible extension to the current curation regarding a certain model should be made after a cross-check with all the BioModels regarding the same paper. As shown in Table 1, there are three papers covering multiple BioModels. In detail, BIOMD0000000780, BIOMD0000000781, and BIOMD0000000782 correspond to one paper [52]; BIOMD0000000793 and BIOMD0000000795 correspond to one paper [58]; and BIOMD0000001037 and BIOMD0000001038 correspond to one paper [64].

Following the validated, extended, and corrected examples in the three BioModels BIOMD0000001023, BIOMD0000000939, and BIOMD0000000839, readers could also go to GitHub (https://github.com/sys-bio/Developing-a-workflow-for-creating-OMEX-files) to cross-check all our successfully reproduced models by comparing the reproduced plots with their original results in the corresponding papers. The original curation from the BioModels Database was stored under the folder original curation inside each BioModel folder, i.e., BIOMD000000XXXX, as a comparison to illustrate our improvement. Figures available on GitHub could also get re-generated via the OMEX files provided on GitHub by a simple type in the Tellurium IDE component “IPython Console”: te.executeCombineArchive(“path to/BIOMD000000XXXX.omex”)

The 50 successfully validated and corrected BioModels curation on GitHub illustrated that our manual workflow described in the section Materials and Methods worked somehow with the current curation and tooling. However, there were still some limitations.

### Non-reproduction due to tooling

The tooling we used was Tellurium with Antimony, RoadRunner, libSBML, libSEDML, and phraSED-ML imported. The successful 50 reproduced models illustrated the advantages of these tools, however, the current tooling also had some limitations. The figures that we successfully reproduced were mostly simulations based on time courses. While more complex figures were not able to be reproduced due to the software and standards used, i.e., figures of bifurcation, 3D plots, etc. For example, we were not able to create an OMEX file following our manual workflow for the phase portraits of Fig 1 and Fig 2 nor the 3D plots of Fig 4 and Fig 5 in the Alharbi paper [64], because the current version of SED-ML does not support it. Therefore, the tooling needs to be improved and advanced in the future for curation.

In addition, it would be good for SED-ML to combine two plots into one in the future, but the current tooling does not make use of this capability. For instance, BIOMD0000000815 has an interesting procedure of cell density change at the beginning, elimination, after the change, and escape. We have extended the curation to cover Fig 7 in the paper [63]. However, the current tooling could only display the procedure in two separate figures as shown in Fig 7.

**Fig 7.**
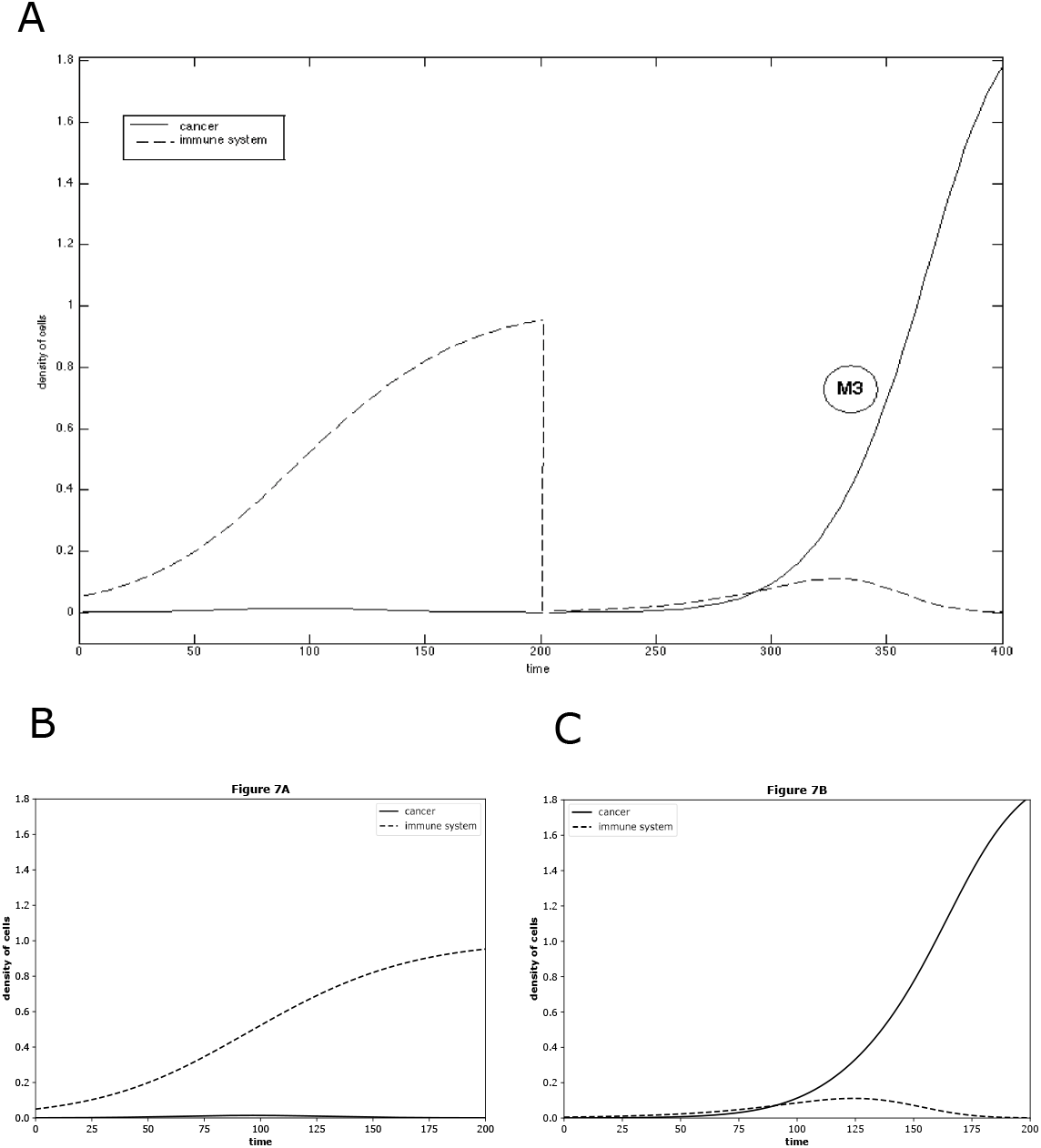
The reproduced results based on BIOMD0000000815. BIOMD0000000815 describes a mathematical model of induced cancer-adaptive immune system competition. (A) is the original result published in the paper [63] as Fig 7. The figure illustrates the evolution of the area of the sarcoma for mice M3, from the beginning, elimination, to the treatment, escape. (B) and (C) indicated the comparable results reproduced by Tellurium.

Furthermore, SED-ML does not support the capacity to insert subplots into one plot nor support bar plots. For instance, in MODEL2002110001 corresponding to the paper [68], Fig 2a has a bar subplot as shown in Fig 8. This can be expressed in SED-ML, but few interpreters yet support these features. However, it is possible that SED-ML can store all the simulations regarding subplots and bar plots in the future, as we successfully reproduced this model by Tellurium and curated the simulation by

**Fig 8.**
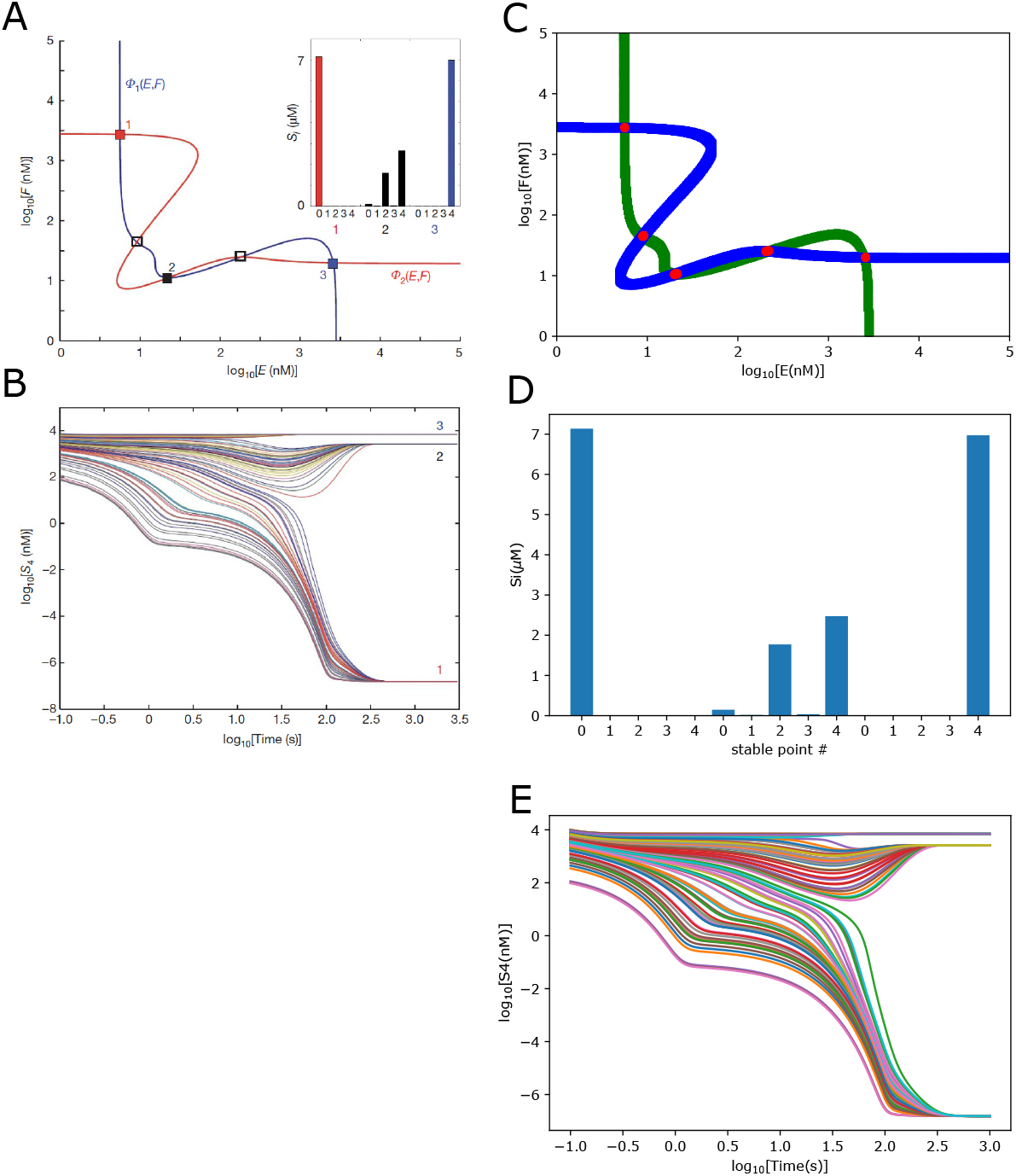
The reproduced results based on MODEL2002110001. MODEL2002110001 describes the unlimited multistability in multisite phosphorylation systems. (A) and (B) are the original results published in the paper [68] as Fig 2. The figure illustrates the Multistability of an *n* = 4 distributive sequential system. (C), (D), and (E) indicated the comparable results reproduced by Tellurium.

Python scripts in BioModels Database (https://www.ebi.ac.uk/biomodels/MODEL2002110001#Overview) and GitHub (https://github.com/SunnyXu/Unlimited_multistability) several years ago.

### Non-reproduction due to paper information

In addition, not all the figures in one paper were reproducible due to errors or a lack of information from the corresponding papers. For instance, some parameters were given incorrectly or even not given. Some figures needed experimental data to fit, while some formulas or functions were not given to reproduce the dynamical behaviors. There are some examples shown in Table 2. See S1 Appendix for more details.

**Table 2.** Non-reproduction examples with certain limitations from papers.

| Limitations | Examples |
| --- | --- |
| non-reproduction due to given parameters | 548, 642, 757, 780, 877, 949, and 984 |
| non-reproduction due to lack of formulae | 005 |
| non-reproduction due to lack of data | 745 and 909 |
| non-reproduction due to lack of related instructions | 953 |

The numbers in the Examples column are the last three digits of the ID from the certain related BioModel.

Therefore, it would be good to provide the data to plot or fit in the paper. For instance, Fig 2B in the BIOMD0000000839 corresponding to the paper [65] shows data that is unavailable, and therefore cannot be added to the plot. See Fig 5C and Fig 5D.

## Discussion

The significance of reproducibility for scientific research has grown substantially over time. Systematic curation has similarly become increasingly important in effectively utilizing published data. We utilized systems biology standards and supportive tools to analyze a selection of models from the BioModels Database. Our work cross-checked 50 BioModels and corrected/extended some of them as necessary. The reproducibility of the 50 models allowed us to develop a manual workflow. There are over a thousand curated models in the BioModels Database, and there are also other models to reproduce beyond this Database. Therefore, it would be impossible to validate, correct, or extend the whole curation manually. However, our manual workflow could be a start to help us achieve a possible automatic workflow to generate an automatic online platform in the future, i.e., https://biosimulations.org and www.reproducibilityportal.org, eventually to help users more easily curate models, or expand models from public repositories. Our colleagues are currently working on another article for the automation to “Read and modify the SED-ML file” for a thousand BioModels (https://github.com/sys-bio/temp-biomodels/tree/main/final). We plan to expand the workflow to cover the whole BioModels Database and beyond automatically in the future. The successfully 50 reproduced models can be a test case for a potential future automatic curation procedure.

We also examined the current limitations and potential improvements in curation practices, standards, and toolsets. In the BioModels Database, the SED-ML is not always present, and even when it is, is not always able to reproduce the figures shown in the corresponding papers. In addition, there is a potential to extend the curation with more results, which means some of the current entries could be extended.

The tooling we used was Tellurium, with Antimony RoadRunner, libSBML, libSEDML, and phraSED-ML imported. The successful 50 reproduced models illustrated the advantages of these tools, however, the current tooling had some limitations. The figures that we successfully reproduced were mostly simulations based on time courses, while more complex figures were not able to be curated as OMEX files, i.e., figures of bifurcation, 3D plots, or bar graphs. Therefore, the tooling needs to be improved in the future. It would also be good for more interpreters to implement certain advanced SED-ML features such as combined subplots. The current manual workflow considered only SBML files as model files in the BioModels Database in the format of SBML. However, it is possible to include other modeling standards in the future, i.e., CellML and NeuroML. As known, models can be encoded using SBML, CellML, or the NeuroML files, and archives containing models in any format could be distributed with the extension of .omex [12]. In addition, once a collection of models is annotated and made available as OMEX archives, the libOmexMeta includes the annotation support for SBML and other modeling languages, such as CellML [69].

In addition, not all the figures in one paper were reproducible due to a lack of information provided in the paper [5, 70]. For instance, some parameters were given incorrectly or not given. Some figures needed experimental data to fit, while some formulas or functions were not given to reproduce the dynamical behaviors. Therefore, it would be good for the authors to publish papers with corresponding data [71, 72]. It would also be good for authors and/or curators to store all the information regarding the publications, i.e., models by SBML files, simulation by SED-ML files, related experimental data, code in repertoire, and possible attachment of the corresponding article.

## Acknowledgments

J.X. appreciates John H. Gennari and Herbert M. Sauro’s comments on the early stage of the manuscript. J.X. also thanks Frank T. Bergmann for his assistance in using libSEDML. J.X. is grateful for the valuable comments from reviewers which have improved this work.

## Competing Interests

The authors have declared that no competing interests exist.

## Data availability statement

The reproduced models and code for this project are fully available on GitHub (https://github.com/sys-bio/Developing-a-workflow-for-creating-OMEX-files) without restriction.

## Funding

JX and LS are grateful to the generous support from National Institute of Biomedical Imaging and Bioengineering of the National Institutes of Health (https://www.nibib.nih.gov/) under award number P41EB023912. The funders had no role in study design, data collection and analysis, decision to publish, or preparation of the manuscript.

## Author contributions

Conceptualization: Jin Xu, Lucian Smith; Data curation: Jin Xu; Formal analysis: Jin Xu; Investigation: Jin Xu; Methodology: Jin Xu, Lucian Smith; Project administration: Jin Xu; Resources: Jin Xu; Software: Jin Xu; Supervision: Jin Xu, Lucian Smith Validation: Jin Xu; Visualization: Jin Xu; Writing – original draft: Jin Xu; Writing – review & editing: Lucian Smith.

## Supporting information

**S1 Table. Finding parameter values in papers**.

We mostly referred to the SBML files for the model information. However, we also referred to the corresponding papers to validate, correct, and extend the models. Most parameter information was provided in the figure titles, however, we also found some valuable information in the main text and tables in the paper. Therefore, we suggest checking parameters following the order of SBML files, figure titles, text content, and tables in the papers.

**Table S1.** Finding parameter values in papers.

| BioModel ID | Figure title | Text | Table | BioModel ID | Figure title | Text | Table |
| --- | --- | --- | --- | --- | --- | --- | --- |
| BIOMD0000000003 |  |  |  | BIOMD0000000850 | X |  |  |
| BIOMD0000000005 | X | X |  | BIOMD0000000877 | X | X |  |
| BIOMD0000000010 |  |  |  | BIOMD0000000894 |  |  |  |
| BIOMD0000000079 | X |  |  | BIOMD0000000909 | X |  |  |
| BIOMD0000000548 | X |  |  | BIOMD0000000911 |  |  |  |
| BIOMD0000000552 | X |  |  | BIOMD0000000916 | X |  |  |
| BIOMD0000000555 |  |  |  | BIOMD0000000930 | X |  |  |
| BIOMD0000000618 |  |  |  | BIOMD0000000932 |  |  |  |
| BIOMD0000000642 |  |  |  | BIOMD0000000933 |  |  |  |
| BIOMD0000000667 |  |  |  | BIOMD0000000939 |  | X |  |
| BIOMD0000000671 |  |  |  | BIOMD0000000947 |  |  |  |
| BIOMD0000000704 | X |  |  | BIOMD0000000948 |  |  |  |
| BIOMD0000000712 |  |  |  | BIOMD0000000949 | X |  | X |
| BIOMD0000000720 | X |  |  | BIOMD0000000953 |  |  |  |
| BIOMD0000000745 |  | X |  | BIOMD0000000964 | X |  |  |
| BIOMD0000000757 | X | X |  | BIOMD0000000967 |  |  |  |
| BIOMD0000000780 | X | X |  | BIOMD0000000970 |  |  |  |
| BIOMD0000000781 | X | X |  | BIOMD0000000984 |  | X |  |
| BIOMD0000000782 | X | X |  | BIOMD0000000986 |  |  |  |
| BIOMD0000000785 | X |  |  | BIOMD0000001004 |  |  |  |
| BIOMD0000000793 |  |  |  | BIOMD0000001006 |  |  |  |
| BIOMD0000000795 |  |  |  | BIOMD0000001023 |  |  |  |
| BIOMD0000000799 | X |  |  | BIOMD0000001026 | X |  |  |
| BIOMD0000000815 | X |  |  | BIOMD0000001037 |  |  |  |
| BIOMD0000000839 |  |  |  | BIOMD0000001038 |  |  |  |

The BioModel ID, Figure title, Text, and Table columns are for the model IDs in the BioModels Database, the sentences in the figure title, the text content in the paper, and the tables in the paper.

**S1 Appendix. Non-reproduction examples with certain limitations from papers**.

For BioModels 548, 642, 757, 780, 877, 949, and 984, there were non-reproductions due to given parameters. We were able to reproduce some of the figures in the paper, however, were not able to reproduce the other relevant figures by only adjusting the corresponding parameters provided. In detail, we were able to reproduce Fig 3 in BioModel 548, but were not able to reproduce Fig 2 by only adjusting the parameter *m* from 25 to 10. We were able to reproduce Fig 1 in the BioModel 642 with parameter *pi* = 0 but were not able to reproduce Fig 2 and Fig 3 with parameters *pi* = 0.3 or *pi* = 0.6. For BioModel 757, we successfully reproduced Fig 1, but couldn’t reproduce Fig 2 and Fig 3 with the parameters given. For BioModel 780, we were able to reproduce Fig 6, but couldn’t reproduce Fig 7 and Fig 8 with the parameters given. For BioModel 877, we were able to reproduce Fig 1 and Fig 2, but couldn’t reproduce Fig 3 and Fig 4 with the parameters given. For BioModel 949, we were able to reproduce Fig 2, but couldn’t reproduce Fig 3 with the parameters given. For BioModel 984, we were able to reproduce Fig 3, but couldn’t reproduce Fig 4 and Fig 5 with two small values of parameter *k*.

For BioModel 005, Fig 3c was not reproducible due to the lack of formula of *k*_6_. For BioModel 745, there were no data of tumor volume data given to reproduce Fig 5 and Fig 6. For BioModel 909, the data in Fig 11 and Fig 12 were not given. For BioModel 953, we were able to reproduce Fig 7B, but there were no clues regarding how to reproduce Fig 7C.

## Notes

### Competing Interest Statement

The authors have declared no competing interest.

### Summary of Updates

Some sentences in the sections of Introduction, Methods and Materials, and Discussion are rephrased to make everything clearer.

